# Emergence and antigenic characterisation of influenza A(H3N2) viruses with hemagglutinin substitutions N158K and K189R during the 2024/25 influenza season

**DOI:** 10.64898/2026.02.10.704996

**Authors:** Björn Koel, Alexander MP Byrne, Sam Turner, Sarah James, Ruth Harvey, Monica Galiano, Christine Carr, Pascal Lexmond, Mark Pronk, Ruonan Liang, Geert-Jan Boons, Robert P. de Vries, Dirk Eggink, Nicola Lewis, Derek Smith, Adam Meijer, Ron Fouchier

## Abstract

**Background:** Seasonal human influenza viruses can escape from antibody-mediated neutralization when amino acid changes occur in the hemagglutinin protein. Routine surveillance identified circulation of an A(H3N2) virus variant in the Netherlands with amino acid substitutions at hemagglutinin positions 158 and 189. These amino acid positions were previously responsible for antigenic change of influenza A(H3N2) viruses and potentially lead to escape of this variant from vaccine-mediated immunity.

**Aim:** To characterize the emergence and antigenic properties of N158K and K189R double substitution virus variants.

**Methods:** We analyzed the geographical and temporal dynamics of the double-substitution variant using a phylogeographic approach and used hemagglutination inhibition assays and antigenic cartography methods to map its antigenic properties.

**Results:** A(H3N2) viruses carrying K189R were first detected in Guatemala in June 2024, before subsequently gaining the N158K substitution, which was intially detected in Colombia in November 2024, followed by detection in the Netherlands in December 2024. However, detections within Europe remained almost entirely confined to the Netherlands. The proportion of viruses carrying the N158K and K189R substitutions increased to 16% - 24% per collection week of sequenced Dutch viruses during the peak of the epidemic of the 2024-2025 respiratory season. Antigenic characterization of viruses with N158K and K189R substitutions indicated that these are antigenically distinct from the A(H3N2) components of 2025-2026 Northern Hemisphere vaccines, showing 8–192-fold reduction in hemagglutination inhibition titers with antisera against the vaccine strain compared to antisera against the homologous virus.

**Conclusions:** Influenza A(H3N2) viruses with N158K and K189R escaped recognition by antibodies raised against the 2024-2025 and 2025/2026 Northern Hemipshere vaccine strains in hemagglutination inhibition assays. These variants circulated widely in the Netherlands during the 2024-2025 influenza season, raising concerns about reduced vaccine-mediated protection if such variants would spread more broadly during 2025-2026 Northern Hemipshere season.

## Introduction

Influenza viruses of the A(H3N2) subtype have been circulating in the human population since 1968 and are a major cause of annual influenza epidemics. Antibodies targeting the hemagglutinin (HA) surface protein can effectively neutralize influenza viruses and are elicited in response to infection or vaccination. However, continuous changes in the HA amino acid composition results in evasion of neutralizing antibodies in the population through antigenic drift. Major antigenic differences between A(H3N2) viruses have historically been caused by amino acid substitutions adjacent to the HA receptor binding site (RBS) ^1–3^, highlighting the importance of monitoring amino acid changes at these positions.

The WHO Global Influenza Surveillance and Response System (GISRS) is a global network of National Influenza Centers (NICs) that monitor influenza viruses. Representative influenza virus positive clinical specimens and/or virus isolates, and data from genetic and phenotypic analyses performed at the NICs, are shared with the WHO Collaborating Centers to inform the WHO recommendation on the composition of influenza vaccines. As part of routine surveillance activities coordinated through the Dutch NIC, a genetic A(H3N2) virus variant carrying HA amino acid substitutions N158K and K189R was identified, starting early in the 2024/2025 influenza season. Both amino acid positions are immediately adjacent to the HA RBS and substitutions at these positions have previously been responsible for large antigenic changes during A(H3N2) virus evolution^3^, raising concerns about the potential spread of this variant and its ability to evade antibodies elicited by the seasonal influenza vaccine. Here, we describe the emergence and spatiotemporal distribution of this A(H3N2) double-substitution variant using phylogeographic methods, as well as the results of antigenic characterization.

## Methods

### Phylogenetic and phylogeographic analysis

A dataset consisting of influenza A virus HA sequences of the H3N2 subtype was constructed using sequences available in the GISAID EpiFlu Database (https://www.gisaid.org) as of 24th March 2025. This data was curated to focus on sequences with collection dates after 1st February 2024, representing the current and prior Northern Hemisphere Influenza seasons. Additionally, representative sequences with collection dates prior to 1st February 2024, were included to provide genetic context and outliers were removed manually. The final sequence dataset is available via the GISAID identifier: EPI_SET_250402wk (Supplemental Table 1). The sequences were aligned using Nextclade version 3.3.0^1^ with the ‘nextstrain/flu/h3n2/ha’ dataset. Time-resolved maximum-likelihood phylogenies were inferred using IQ-Tree2^2^ and TreeTime^4^ as implemented in the Augur toolkit^5^ using a fixed clock-rate and clock-rate standard deviation of 3.82x10^-3^ and 7.64x10^-4^ nucleotide substitutions per year, respectively, obtained from the seasonal influenza build available on Nextstrain^6^. Ancestral sequence inference and translation, as well as discrete trait analysis to assess viral dispersal were also performed using the Augur toolkit. Final annotated phylogenies were analysed and visualised using Auspice within the Nextstrain platform.

### Sample collection

The Dutch National Influenza Centre (NIC), operating from the Erasmus Medical Centre (EMC), the National Institute for Public Health and the Environment (RIVM), and Nivel, obtains clinical specimens from three sources; i) sentinel general practitioners submit combined nasopharyngeal/oropharyngeal swab specimens from patients presenting with influenza like illness or another acute respiratory infection, ii) Dutch hospital and peripheral diagnostic laboratories submit influenza positive samples from mainly hospitalized patients and outpatients, and iii) patients with a self-reported acute respiratory infection submit a self-collected nose/throat swab specimen through participatory community surveillance (“Infectieradar”)^4^.

### Genetic characterization

Dutch specimens positive for influenza A virus were genetically characterized by subtype-specific PCR and/or nanopore sequencing. Briefly, viral RNA was isolated using the Roche High Pure RNA Isolation kit or MagNA Pure 96 DNA and Viral NA Small Volume kit (Roche, Basel, Switzerland) according to the manufacturer’s instructions and amplified using a universal eight-segment PCR using UNI primers^5^. The PCR products were purified, pooled, and further processed using the Nanopore Ligation Sequencing Kit (Oxford Nanopore, Oxford, the United Kingdom). Subsequently, the preprocessed PCR products were sequenced on a GridION Flow Cell (R10.4.1) platform (Oxford Nanopore) and consensus sequences were obtained using CLC Genomic Workbench (CLC bio, Cambridge, MA, USA) or ViroConstrictor version 1.4^6^. Viruses with relevant genetic signatures were selected and isolated in modified Madin-Darby canine kidney (hCK) cells^7^ or mixed Madin-Darby canine kidney (MDCK)-2,6-sialyltransferase (SIAT)/MDCK-I cells^8^ for subsequent antigenic characterization. The final sequence dataset generated from the Netherlands is available via the GISAID identifier: EPI_SET_251001mv (Supplemental Table 2).

### Antigenic characterization

Ferret antisera were prepared by intranasal inoculation with 500µL virus stock. Antisera were collected 14 days after inoculation. The generation of ferret antisera at Erasmus MC was carried out in accordance with the Dutch legislation for the protection of animals used for scientific purposes (2014, implementing EU Directive 2010/63). The study was approved by the Central Committee on Animal Experiments (CCD), under project license CCD101002115685 and AVD27700202216424. The generation of ferret antisera at the Francis Crick Institute was carried out in accordance with UK legislation for the use of animals under the Animals (Scientific Procedures) Act 1986 under project license PP4282545.

Due to reduced agglutination of turkey erythrocytes by single- and double-substitution variants, hemagglutination inhibition (HI) assays were performed using guinea pig erythrocytes (Francis Crick Institute) or using glycoengineered turkey erythrocytes (EMC). For the latter, turkey erythrocytes were enzymatically modified to install extended LacNAc moieties terminating in α2,6-sialosides, as described previously^9^ and kept in a 50% solution in DPBS (Sigma-Aldrich, Zwijndrecht, The Netherlands).

HI assays were performed as described previously^10^. Briefly, ferret antisera were pre-treated overnight with Vibrio Cholera Neuraminidase at 37°C followed by inactivation at 56°C for one hour. Two-fold serial dilutions, starting at a 1/20 dilution, of the pre-treated sera were mixed with 4 HA units of diluted virus stock in 25µL. Subsequently, 25 µL 1% erythrocytes was added and the mixture was incubated for one hour at 4°C before reading agglutination patterns. The HI titer is expressed as the reciprocal of the highest antiserum dilution that completely inhibited agglutination of erythrocytes. Analysis of antigenic properties was supported by antigenic cartography methods which facilitate the quantification and visualization of HI assay data^11^. In an antigenic map, the distances between antigens and antisera represent antigenic distance as measured by the HI assay, with distances inversely related to the log2 HI titer. As each antigen is tested against multiple antisera and vice versa, the measurements can be used to determine the position of antigens and antisera in the map. Antigenic map construction was performed using the Racmacs R package version 1.2.9^12^.

## Results

### Emergence and dispersal of A(H3N2) J.2 influenza viruses with the N158K K189R double-substitution

Since the 1st February 2024, the 2a.3a.1 clade has been the predominant A(H3N2) clade circulating globally, and the HA gene of these viruses has undergone further diversification into multiple subclades (J.1-J.4), of which the J.2 subclade has been most prevalent^7^ (Figure 1A). Within the J.2 subclade, a number of amino acid substitutions have emerged in the HA protein, including those at positions 158 and 189. The ancestral amino acid identified at position 158 was an asparagine, however, substitutions including lysine, histidine, serine and threonine have been observed with the asparagine to lysine (N158K) substitution being the most common (Figure 1B). At position 189, substitutions from lysine to asparagine, arginine and serine have been observed, but the lysine to arginine (K189R) substitution has been most prevalent (Figure 1C).

**Figure 1.**
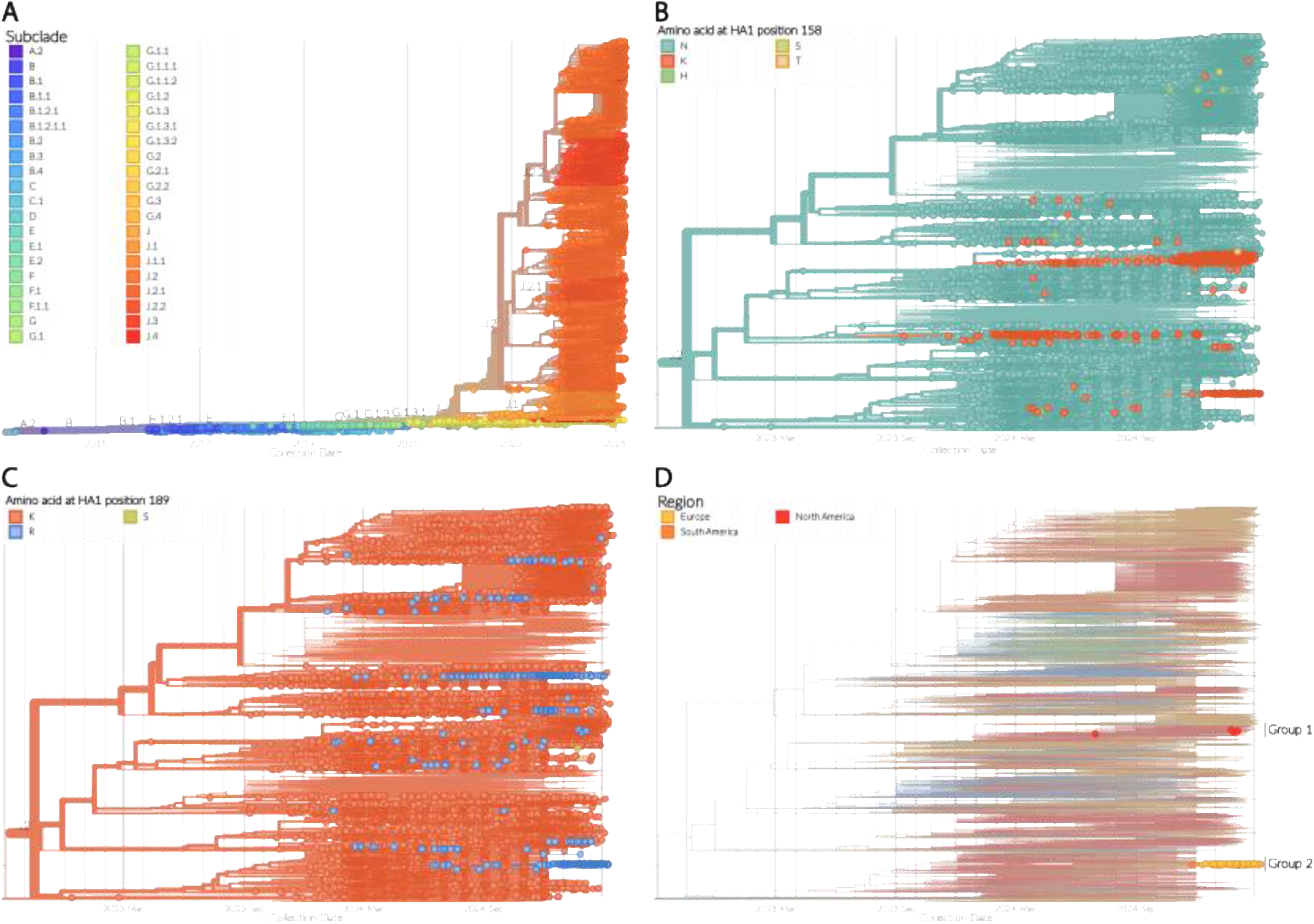
Phylogenetic analysis of human A(H3N2) influenza A viruses. **A)** Time-resolved maximum-likelihood phylogenetic tree of the HA gene segment of contemporary human A(H3N2) influenza A viruses constructed using the specimen collection date. The tip shapes are coloured according to subclade, and the branches within the tree where the subclades diverge are annotated. **B-D)** Magnified images of the phylogenetic tree shown in **A**, focusing on the J.2 A(H3N2) subclade. For **B** and **C**, the tip shapes are coloured according to the amino acid found at HA1 positions 158 and 189, respectively. For **D**, only the sequences which contain both N158K and K189R substitutions are highlighted, with the tip shapes coloured according to region of origin. The two distinct groups of A(H3N2) J.2 sequences containing both of these substitutions are annotated.

In addition to singular substitutions, there are also J.2 sequences that contain both the N158K and K189R substitutions, which form two distinct groups within the phylogenetic tree (Figure 1D), showing these substitions in different genetic backgrounds. The first group consists of four sequences from the United States with collection dates ranging from the 1st July 2024 to 4th February 2025 (Figure 1D Group 1). The sequences in this group do not form a monophyletic group and sit within a larger cluster of sequences which all contain the N158K substitution (Figure S1) suggesting the sporadic occurrence of the N158K and K189R double substitutions on multiple occasions. The second group of sequences with the N158K and K189R double substitutions consists of 107 sequences from South America (N=1; Colombia), North America (N=10; the United States) and Europe (N=96; the Netherlands, N=95 and Sweden, N=1), with collection dates ranging from 26th November 2024 through to 11th March 2025 based on the timeframe evaluated. These sequences sit within a wider phylogenetic cluster containing the K189R substitution and suggest that the N158K substitution (along with a S49N substitution in HA2) emerged on a singular occasion with subsequent spread of the double mutant (Figure 2).

**Figure 2.**
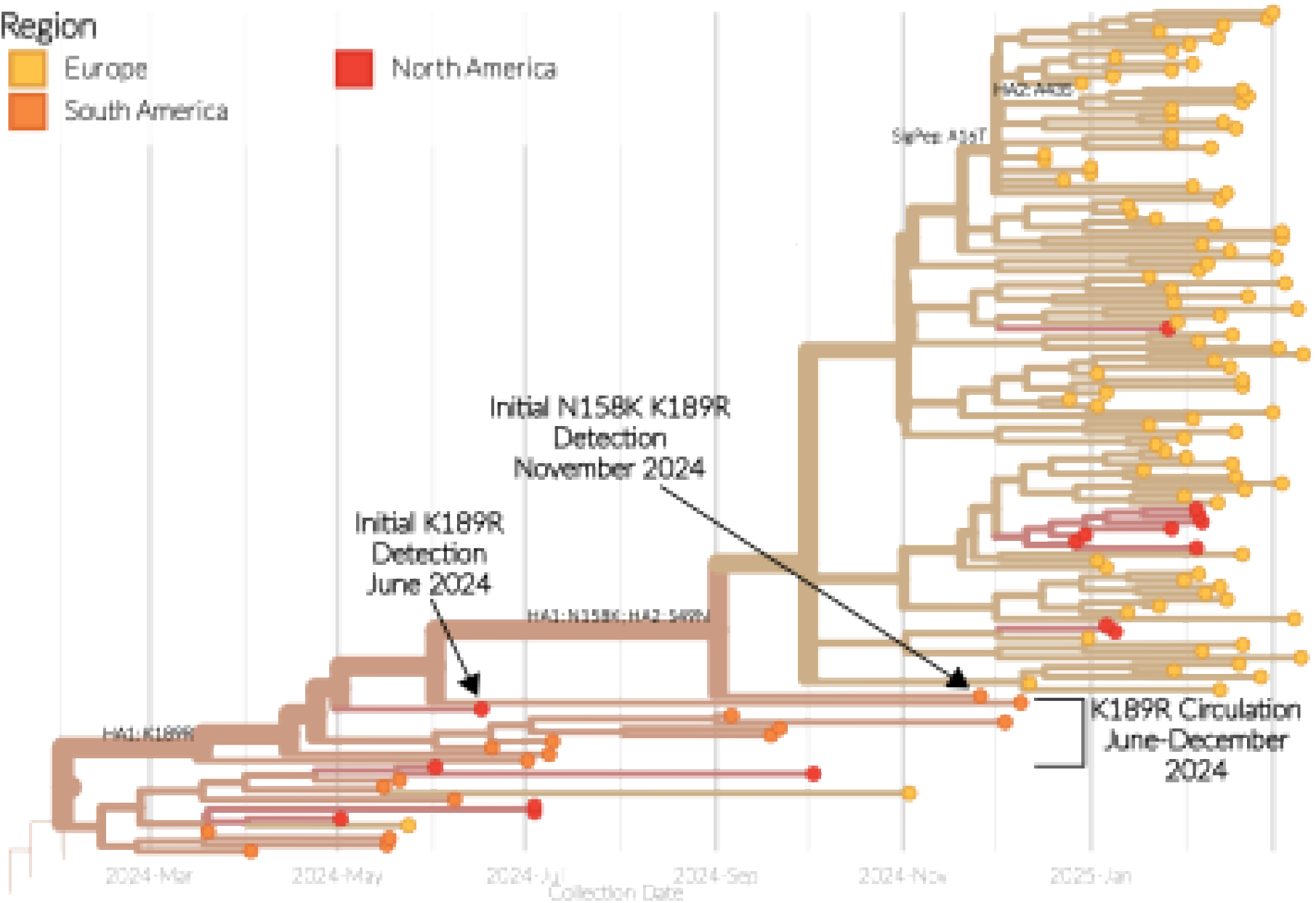
Emergence of the A(H3N2) J.2 viruses with N158K K189R double-substitution. Magnified images of the time-resolved maximum-likelihood phylogenetic tree of the HA gene segment from Figure 1A demonstrating the emergence of the K189R and N158K substitutions. Key dates and events are annotated. Tip shapes and branches are coloured according to the region of origin.

Using time-resolved phylogenies and discrete trait analysis, it is possible to determine the timeline and dispersal of these viruses. The clade containing the K189R substitution (which subsequently obtained N158K also) was initially detected in a virus on the 17th June 2024 in Central America (Guatemala), before detections on the 20th Jun 2024 in South America (Brazil), where it continued to be detected until December 2024 (Figure 2). All of the viruses containing the K189R substitution share a common ancestor dating back to March 2025 (confidence interval: February – April 2024), and were inferred to have emerged in South America, which suggests undetected circulation prior to the detections in June 2024.

The N158K substitution was first detected on 26th November 2024, again in South America (Colombia), before being detected in Europe (the Netherlands) on 12th December 2024. After Europe, there were subsequent detections in North America (the United States) between 27th December 2024 and 6th February 2025. Based on the phylogeny, all of the N158K K189R double-substitution viruses share a common ancestor dating back to September 2024 (confidence interval: June – November 2024), which was inferred to also have emerged in South America. There was then a single introduction into Europe, but three discrete and separate introductions from Europe into North America.

### Circulation of the N158K K189R double-substitution variant in the Netherlands

The N158K K189R double-substitution variant was first detected in the Netherlands in week 50 of 2024, in a clinical specimen submitted to the NIC by a sentinel general practitioner (Figure 3). From week 50 of 2024 until week 10 of 2025, a total of 594 A(H3N2) virus sequences from the Netherlands were submitted to GISAID by the NIC, 87 of which contained the N158K K189R double-substitution. Following the initial detection, an increasing number and proportion of N158K K189R double-substitution variants was observed (Figure 3).

**Figure 3.**
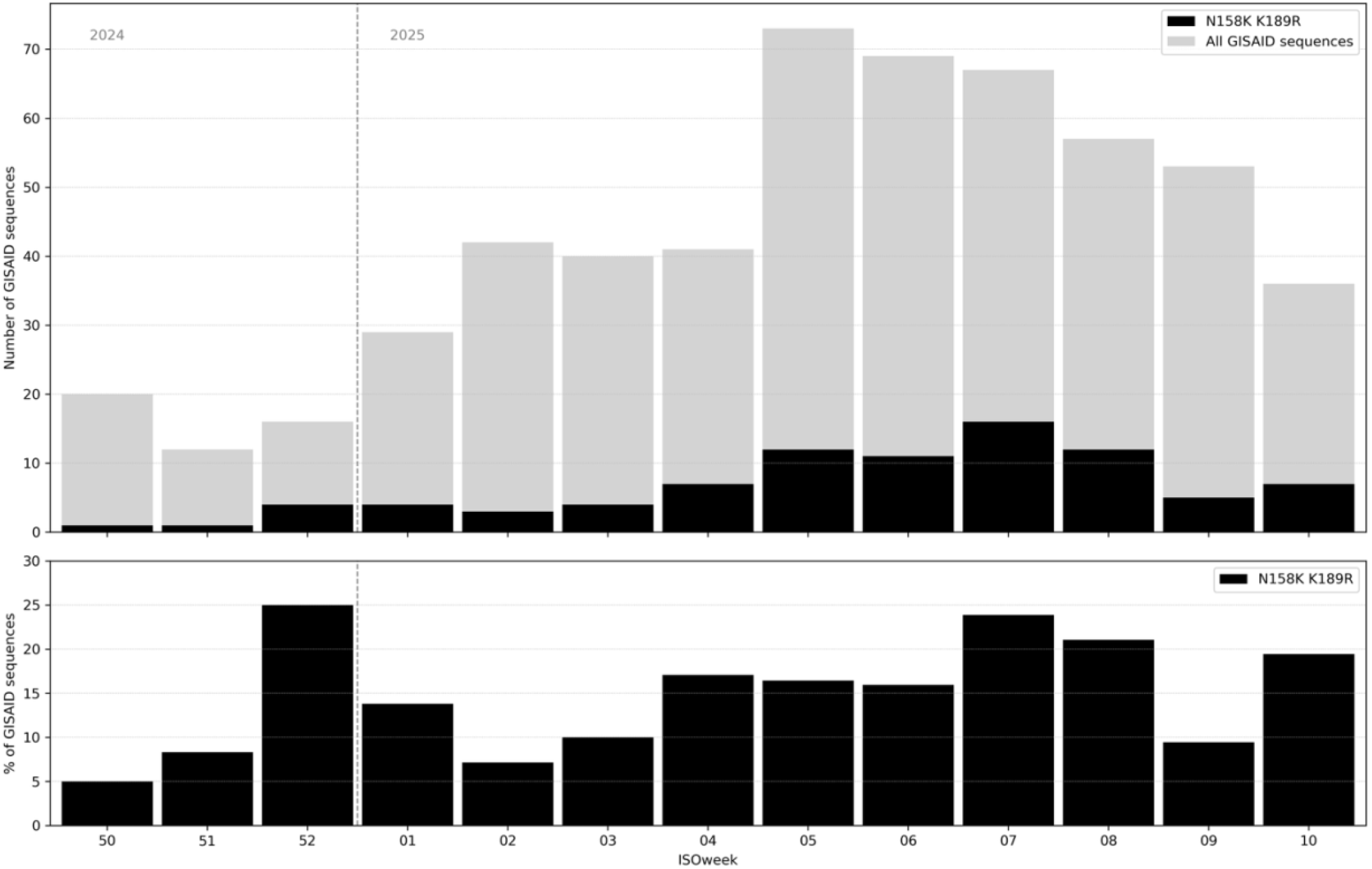
Circulation of viruses with the N158K K189R double-substitution in the Netherlands, weeks 40 of 2024 through week 10 of 2025. Number and proportion of total sequences of N158K and K189R double-substitution viruses uploaded to GISAID, by collection week, with original specimen collection date from ISOweek 40 2024 (starting Monday December 9^th^) until ISOweek 10 of 2025 (starting Monday March 3^rd^). No double-substitution viruses were identified in the Netherlands prior to week 50.

Double-substitution variants were detected across multiple regions of the Netherlands and in all three national surveillance systems: sentinel general practitioners, diagnostic laboratories, and participatory community surveillance. The weekly count and proportion of the double-substitution variant peaked between weeks 5 and 8 of 2025, coinciding with the peak of the influenza epidemic in the Netherlands. During this period, 11 to 16 detections were observed per week. The double-substitution variant accounted for 16% to 24% of all submitted sequences per week, based on collection date. The proportion of double-substitution variants was also high in week 52 of 2024 (25%, 4 detections among 16 submitted sequences), but because of the relatively low number of virus detections that week the estimated proportion should be interpreted with caution. Single-substitution variants N158K and K189R were each detected twice; N158K in weeks 41 and 42 of 2025, and K189R in weeks 7 and 8 of 2025.

### Antigenic characterization of N158K K189R double-substitution variants

To determine the antigenic effect of N158K and K189R we characterized viruses carrying these amino acid substitutions, both individually and in combination, using hemagglutination inhibition (HI) assays. Additionally, contemporary variants with T135K or T135A substitutions, which disrupt the N-linked glycosylation motif at position 133 and potentially alter the antigenic properties, were included in the HI assay. The antigenic properties of each of these variants was mapped in relation to the A(H3N2) vaccine strains for the Northern Hemisphere (NH) 2024– 2025 season (A/Thailand/8/2022 egg-based, A/Massachusetts/18/2022 cell-based) and NH 2025–2026 vaccine strains (A/District of Columbia/27/2023 cell-based, A/Croatia/10136RV/2022 egg-based), which are identical to the strains recommended for the Southern Hemisphere 2025 season. To account for differences in isolation methods used by the National Institute for Public Health and the Environment and Erasmus MC (see methods section), a subset of 5 double-substitution viruses collected by both institutes during the 2024–2025 season was tested in the HI assay. The antigenic properties of all viruses were analysed using a panel of ferret antisera. As viruses carrying substitution T135K and N158K agglutinated turkey erythrocytes poorly, glycan-modified turkey erythrocytes and guinea pig erythrocytes were used to facilitate reliable antigenic characterisation.

In the HI assay using glycan-modified erythrocytes, viruses with substitution N158K showed antigenic divergence from the NH 2024-2025 and NH 2025-2026 recommended vaccine strain for A(H3N2) virus (Figure 4A and Supplemental Table 3). Viruses with substitution N158K had 1.5–6-fold reduced titers to the antiserum raised against A/Thailand/8/2022 compared to the homologous virus, 4–8 fold reduced titers to the A/District of Colombia/27/2023 antiserum, and 6–24-fold reduced titers to the A/Croatia/10136RV/2022 antiserum. Viruses carrying substitution K189R were antigenically distinct from A/Thailand/8/2022 (16-24-fold reduction compared to homologous), but not from A/District of Colombia/27/2023 and A/Croatia/10136RV/2022 (<4-fold compared to homologous). All five Dutch isolates with double substitution N158K K189R were antigenically distinct from the NH 2024-2025 and NH 2025-2026 vaccine strains, showing markedly reduced titers of 8–192-fold compared to homologous titers. Two Dutch isolates, one with the T135A S145N double-substitution and one without either of these substitutions, were antigenically similar to A/Thailand/8/2022. In contrast, other viruses with either T135K or S145N, including the NH 2025–2026 recommended vaccine strains were antigenically different from A/Thailand/8/2022, with 6–12-fold reduced HI titers.

**Figure 4.**
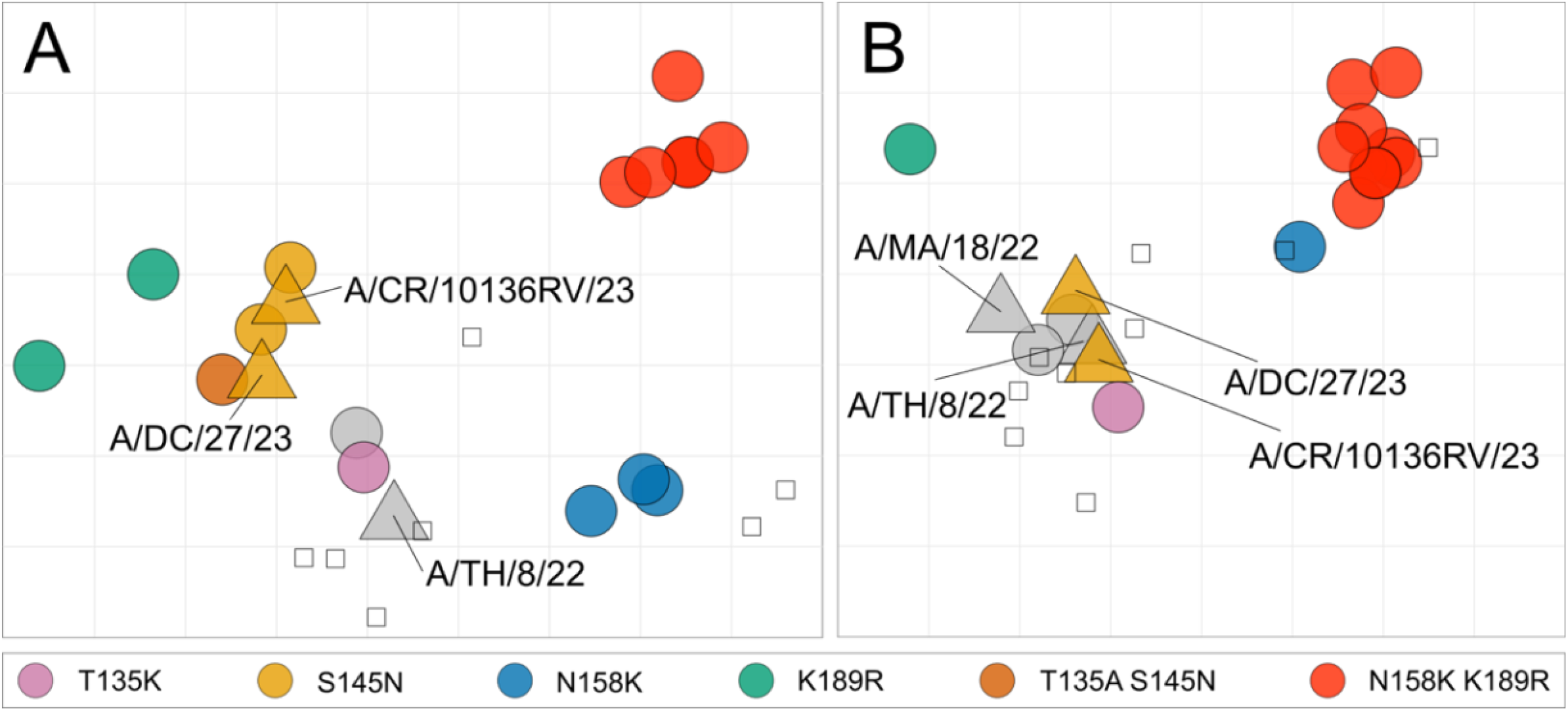
Antigenic maps of recent A(H3N2) viruses. The position of vaccine strains is indicated as filled triangles; all other viruses as filled circles. Viruses are color-coded by HA amino acid substitutions: T135K (purple), S145N (orange), N158K (blue), K189R (green), T135A + S145N (brown), and N158K + K189R (red). Other antigens are indicated in grey. Open squares indicate the position of antisera. Each square in the antigenic map grid corresponds to a two-fold difference in the HI assay titer. Abbreviations used in the figure are as follows: CR, Croatia; DC, District of Columbia; MA, Massachusetts; TH, Thailand. **A)** Antigenic map based on HI assay data from Erasmus MC using glyco-modified turkey erythrocytes. **B)** Antigenic map based on HI assay data from the Francis Crick institute using guinea pig erythrocytes.

Antigenic characterisation using guinea pig erythrocytes in the HI assay generally showed smaller antigenic differences compared to those found in the HI assay using glycan-modified turkey erythrocytes. A/Slovenia/49/2024 and A/Switzerland/47775/2024, that have the N158K or K189R substitution respectively, were antigenically distinct from recent and recommended vaccine strains and showed 2–16-fold reduced titers to antisera raised against A/Thailand/08/2022, A/District of Columbia/27/2023, and A/Croatia/10136RV/2023 compared to homologous. Consistent with the glycan-modified erythrocyte HI assay, viruses with the N158K K189R double substitution were antigenically most distinct from the NH 2024-2025 vaccine strain and NH 2025-2026 vaccine strains, with ≥8-to ≥32-fold reduced titers compared to homologous and many values below the detection limit of the HI assay at the starting dilution used. Viruses with substitutions T135K and S145N were antigenically similar to the NH 2024-2025 vaccine strains, with <4-fold reduced titers compared to homologous (Figure 4B and Supplemental Table 4).

## Discussion

Growing insights into the molecular determinants underlying antigenic variation of influenza viruses have improved the feasibility of sequence-based surveillance. High-throughput screening of circulating influenza viruses based on sequence data facilitates early detection of emerging genetic variants, including potentially antigenically distinct viruses, which can subsequently be characterized in phenotypic assays. This enables efficient and timely insights into potential mismatches between circulating viruses and current vaccine strains. We identified the emergence of A(H3N2) viruses with an N158K K189R double-substitution expected to be antigenically different from 2024-2025 and 2025-2026 Northern Hemisphere vaccine strains, which was later confirmed in HI assays.

Phylogeographic analysis of the A(H3N2) double-substitution variants demonstrated that they emerged by sequential accumulation of firstly the K189R substitution and then the N158K substitution. These variants were first detected in South America, before detection in Europe and North America. Since the end of our study period (24th March 2025), and at the time of writing (2nd January 2026), this double-substitution variant has been detected elsewhere in Europe (Belgium, Denmark, France, Germany, Norway, Portugal, Spain and the United Kingdom), as well as in Asia (Cambodia, Hong Kong and Japan), Africa (South Africa), North America (Canada, Costa Rica and Panama), South America (Bolivia, Brazil, Chile, Ecuador, French Guiana and Peru) and Ocenia (Australia) with the most recent detection occurring in Bolivia in mid-December 2025. It is also of note that whilst detections of these double-substitution viruses waned during the middle of 2025, they have exhibited a resurgence in late 2025, particularly in South America, particularly Brazil and Chile. Interestingly, the emergence of the N158K K189R double-substitution variant has occurred twice within distinct genetic lineages J.2 subclade of A(H3N2) viruses during the NH 2024-2025 influenza season. However, one of these emergences was geographically restricted to North America, whilst the other was not. The genetic and antigenic mechanisms that may have led to this distinct geographic pattern of transmission are not known, and highlight that further investigation is required to elucidate them.

The double-substitution variant exhibited highly reduced HI titers against ferret antisera raised to the NH 2024-2025 A(H3N2) vaccine strains. Importantly, this variant also showed markedly reduced titers to ferret antisera raised against the vaccine strains for the NH 2025-2026 influenza season. The currently observed antigenic change due to substitutions at amino acid positions 158 and 189 is consistent with previous observations of amino acid changes at these key positions in HA. Amino acid substitutions at position 158 contributed to major antigenic changes in A(H3N2) viruses at least twice^3^, albeit with different amino acid substitutions at this position (G158E, E158K). Substitutions at position 189 have also been responsible for major antigenic change of A(H3N2) viruses on at least two occasions, including the K189R substitution present in the current double-substitution variant. A similar antigenic change has also been observed by a substitution at the equivalent amino acid position (185) in A(H5N1) clade 2.1 viruses^13^.

Although viruses with single substitutions N158K or K189R were antigenically divergent from the NH 2024-2025 vaccine strains, they were detected infrequently in the Netherlands during the study period. However, both single substitutions were observed more commonly in viruses circulating outside of the Netherlands prior to the 2024-2025 influenza season based on analysis of GISAID data. As the combined antigenic effect of N158K and K189R was larger than those of either single substitution alone, this may have contributed to the emergence of the double-substitution variant in the Netherlands. Moreover, its appearance early in the influenza season, before widespread influenza virus circulation, may have facilitated its establishment in the population during a time when competition with co-circulating viruses was limited.

While the outcomes of antigenic characterization are based on the use of ferret antisera in HI assays, which serve as a proxy for the broader antibody repertoire in the human population, we anticipate that the antigenic change caused by the N158K K189R double-substitution will result in a substantial reduction in vaccine effectiveness if viruses carrying these substitutions were to spread more widely in the human population. The circulation of this double-substitution variant in the Netherlands for several months may indicate the presence of an antigenic space that can be occupied by newly emerging, antigenically distinct A(H3N2) viruses, leaving room for such viruses to further expand in the human population. Although detections of viruses carrying the N158K K189R double-substitution have subsequently declined in Europe and been replaced by subclade K viruses carrying N158D and K189R, that are likewise antigenically distinct from current vaccine strains^14^, N158K K189R double-substitution variants continue to circulate elsewhere. This highlights that antigenically distinct variants may persist or re-emerge in different geographic regions. Continued sequence-based surveillance, complemented by phenotypic characterization, is essential for the early identification of emerging antigenic variants that may reduce vaccine effectiveness and necessitate timely adaptation of vaccination strategies.

## Acknowledgements

We gratefully acknowledge all data contributors, i.e., the Authors and their Originating laboratories responsible for obtaining the specimens, and their Submitting laboratories for generating the genetic sequence and metadata and sharing via the GISAID Initiative, on which this research is based. We thank participating Dutch general practices and clinical diagnostic laboratories and their patients for their collaboration and providing the diagnostic specimens.

This work was supported by core funding from Dutch NIC activities from the Dutch Ministry of Health, Welfare and Sport, by core funding from the Francis Crick Institute from Cancer Research UK (grant no. CC1114), the UK Medical Research Council (grant no. CC1114), the Wellcome Trust (grant no. CC1114) and by the National Institute of Allergy and Infectious Diseases (NIAID) under award R01 AI165692.

## Notes

### Competing Interest Statement

The authors have declared no competing interest.

