## Supplemental Files for "Emergence and antigenic characterisation of influenza A(H3N2) viruses with hemagglutinin substitutions N158K and K189R during the 2024/25 influenza season"

### Supplementary material

This supplementary material is hosted by Eurosurveillance as supporting information alongside the article [Title], on behalf of the authors, who remain responsible for the accuracy and appropriateness of the content. The same standards for ethics, copyright, attributions and permissions as for the article apply. Supplements are not edited by Eurosurveillance and the journal is not responsible for the maintenance of any links or email addresses provided therein.

#### Supplemental Table 1

Data Availability

GISAID Identifier: EPI_SET_250402wk

DOI: https://doi.org/10.55876/gis8.250402wk
All genome sequences and associated metadata in this dataset are published in GISAID's EpiFlu database. To view the contributors of each individual sequence with details such as accession number, Virus name, Collection date, Originating Lab and Submitting Lab and the list of Authors, visit 10.55876/gis8.250107tk

Data Snapshot
EPI_SET 250402wk is composed of 27111 individual viruses.
The collection dates range from 2013-12-06 to 2025-03-18;
Data were collected in 145 countries and territories.

#### Supplemental Table 2

Data Availability

GISAID Identifier: EPI_SET_251001mv

DOI: https://doi.org/10.55876/gis8.251001mv
All genome sequences and associated metadata in this dataset are published in GISAID's EpiFlu database. To view the contributors of each individual sequence with details such as accession number, Virus name, Collection date, Originating Lab and Submitting Lab and the list of Authors, visit EPI_SET_251001mv

Data Snapshot
EPI_SET_251001mv is composed of 594 individual viruses.
The collection dates range from 2024-09-30 to 2025-03-09;
Data were collected in 1 countries and territories.

**Supplemental Table 3. Hemagglutination inhibition assay results of A(H3N2) virus variants using glycan-modified erythrocytes.**

| | Characterizing amino acid substitution(s) | A/NL/00322/2020 | A/NL/00652/2023 | A/TH/8/2022 | A/DC/27/2023 | A/CR/10136RV/2023 | A/NL/01285/2024 | A/NO/423/2024 | A/SL/49/2024 |
|---|---|---|---|---|---|---|---|---|---|
| A/NL/00322/2020 | | **1280** | 80 | 30 | 160 | 20 | <10 | <10 | 160 |
| A/NL/00652/2023 | | 320 | **1280** | 1280 | 1920 | 1280 | 640 | 640 | 480 |
| A/TH/8/2022 | | 1280 | 2560 | **3840** | 5120 | 3840 | 1280 | 1280 | 1280 |
| A/DC/27/2023 | S145N | 80 | 320 | 480 | **1280** | 1280 | 320 | 320 | 320 |
| A/CR/10136RV/2023 | S145N | 160 | 320 | 320 | 640 | **640** | 320 | 320 | 320 |
| A/NL/01285/2024 | K189R | 10 | 160 | 160 | 640 | 320 | **640** | 20 | 80 |
| A/NO/423/2024 | N158K | 640 | 640 | 2560 | 1280 | 640 | 640 | **15360** | 20480 |
| A/SL/49/2024 | N158K | 160 | 320 | 640 | 640 | 240 | 480 | 10240 | **20480** |
| A/NL/10038/2025 | S145N | 80 | 240 | 320 | 640 | 320 | 320 | 320 | 240 |
| A/NL/10067/2025 | S145N | 40 | 320 | 320 | 640 | 960 | 240 | 320 | 320 |
| A/NL/10737/2024 | T135K + S145N | 40 | 320 | 640 | 960 | 640 | 160 | 160 | 480 |
| A/NL/01809/2024 | T135K | 240 | 960 | 1280 | 2560 | 2560 | 480 | 640 | 1280 |
| A/NL/01829/2024 | K189R | 10 | 120 | 240 | 640 | 320 | 320 | <10 | 40 |
| A/NL/01577/2024 | N158K | 120 | 320 | 640 | 640 | 160 | 320 | 5120 | 10240 |
| A/NL/00520/2025 | N158K + K189R | 20 | 80 | 40 | 80 | 40 | 320 | 1280 | 1280 |
| A/NL/02093/2024 | N158K + K189R | 20 | 80 | 40 | 80 | 40 | 320 | 1280 | 1280 |
| A/NL/00024/2025 | N158K + K189R | 30 | 160 | 40 | 80 | 80 | 640 | 1280 | 1280 |
| A/NL/00328/2025 | N158K + K189R | 20 | 120 | 40 | 80 | 60 | 480 | 1280 | 1280 |
| A/NL/10685/2024 | N158K + K189R | * | * | 20 | * | * | 160 | 640 | 640 |
| A/NL/10103/2025 | N158K + K189R | 10 | 80 | 20 | 40 | 40 | 320 | 1280 | 1280 |

CR, Croatia; DC, District of Columbia; NL, Netherlands; NO, Norway; SL, Slovenia; TH, Thailand.

The antisera (columns) are designated according to the virus (rows) used to prepare the antiserum. An asterisk indicates that the virus – antiserum combination was not tested. When the highest concentration of antiserum did not inhibit hemagglutination, the titer is indicated as lower than the lowest dilution tested (e.g. <10). Homologous titers are shown in bold.

**Supplemental Table 4. Hemagglutination inhibition assay results of A(H3N2) virus variants using guinea pig erythrocytes.**

| | Characterizing amino acid substitution(s) | A/TH/08/2022 | A/DC/27/2023 | A/CR/10136RV/2023 | A/NL/10563/2023 | A/LI/216/2023 | A/SL/49/2024 | A/SW/47775/2024 | A/NL/10685/2024 | A/CA/NSVH102423723/2024 |
|---|---|---|---|---|---|---|---|---|---|---|
| A/MA/18/2022 | | 320 | 320 | 320 | 320 | 320 | 160 | 40 | <40 | 160 |
| A/TH/08/2022 | | **320** | 640 | 640 | 320 | 640 | 320 | 160 | 40 | 320 |
| A/DC/27/2023 | S145N | 160 | **320** | 640 | 320 | 320 | 320 | 160 | 40 | 160 |
| A/CR/10136RV/2023 | S145N | 320 | 640 | **1280** | 640 | 640 | 640 | 160 | 80 | 640 |
| A/NL/10563/2023 | S124N | 160 | 320 | 640 | **640** | 320 | 320 | 160 | 40 | 320 |
| A/LI/216/2023 | S124N | 320 | 640 | 640 | 320 | **320** | 160 | 80 | <40 | 320 |
| A/SL/49/2024 | N158K | 40 | 80 | 160 | 80 | 160 | **1280** | 40 | 640 | 80 |
| A/SW/47775/2024 | K189R | <40 | 160 | 80 | 80 | 40 | <40 | **160** | <40 | 80 |
| A/NL/10685/2024 | N158K + K189R | <40 | <40 | <40 | <40 | 80 | 320 | 40 | **640** | <40 |
| A/CA/NSVH/2024 | T135K | 320 | 320 | 320 | 320 | 320 | 320 | 80 | 40 | **1280** |
| A/NL/00520/2025 | N158K + K189R | <40 | <40 | <40 | <40 | 80 | 640 | 40 | 1280 | 40 |
| A/NL/00401/2025 | N158K + K189R | <40 | 40 | 40 | <40 | 80 | 640 | 40 | 1280 | 40 |
| A/NL/00328/2025 | N158K + K189R | <40 | 40 | 80 | 40 | 80 | 640 | 40 | 1280 | 80 |
| A/NL/00077/2025 | N158K + K189R | <40 | <40 | <40 | <40 | 80 | 640 | 40 | 640 | 40 |
| A/NL/00156/2025 | N158K + K189R | <40 | 40 | 40 | <40 | 80 | 640 | 40 | 1280 | 40 |
| A/NL/00024/2025 | N158K + K189R | <40 | 40 | 40 | <40 | 40 | 320 | 40 | 1280 | 40 |
| A/NL/02175/2024 | N158K + K189R | <40 | 40 | 40 | <40 | 80 | 640 | 40 | 1280 | 40 |
| A/NL/02182/2024 | N158K + K189R | <40 | 40 | 40 | <40 | 80 | 640 | 40 | 640 | 40 |
| A/NL/02093/2024 | N158K + K189R | <40 | <40 | <40 | <40 | 40 | 320 | <40 | 640 | <40 |

CA, Catalonia; CR, Croatia; DC, District of Columbia; MA, Massachusetts; LI, Lisboa; NL, Netherlands; SL, Slovenia; SW, Switzerland; TH, Thailand. A/CA/NSVH102423723/2024 is abbrebviated as A/CA/NSVH/2024.

The antisera (columns) are designated according to the virus (rows) used to prepare the antiserum. An asterisk indicates that the virus – antiserum combination was not tested. When the highest concentration of antiserum did not inhibit hemagglutination, the titer is indicated as lower than the lowest dilution tested (e.g. <10). Homologous titers are shown in bold.
